# A complete *Cannabis* chromosome assembly and adaptive admixture for elevated cannabidiol (CBD) content

**DOI:** 10.1101/458083

**Authors:** Christopher J. Grassa, Jonathan P. Wenger, Clemon Dabney, Shane G. Poplawski, S. Timothy Motley, Todd P. Michael, C.J. Schwartz, George D. Weiblen

## Abstract

*Cannabis* has been cultivated for millennia with distinct cultivars providing either fiber and grain or tetrahydrocannabinol. Recent demand for cannabidiol rather than tetrahydrocannabinol has favored the breeding of admixed cultivars with extremely high cannabidiol content. Despite several draft *Cannabis* genomes, the genomic structure of *cannabinoid synthase* loci has remained elusive. A genetic map derived from a tetrahydrocannabinol/cannabidiol segregating population and a complete chromosome assembly from a high-cannabidiol cultivar together resolve the linkage of *cannabidiolic* and *tetrahydrocannabinolic acid synthase* gene clusters which are associated with transposable elements. High-cannabidiol cultivars appear to have been generated by integrating hemp-type *cannabidiolic acid synthase* gene clusters into a background of marijuana-type cannabis. Quantitative trait locus mapping suggests that overall drug potency, however, is associated with other genomic regions needing additional study.

**Resources available online at: http://cannabisgenome.org**

**Summary:** A complete chromosome assembly and an ultra-high-density linkage map together identify the genetic mechanism responsible for the ratio of tetrahydrocannabinol (THC) to cannabidiol (CBD) in Cannabis cultivars, allowing paradigms for the evolution and inheritance of drug potency to be evaluated.

## Main Text

THCA (delta-9-tetrahydrocannabinolic acid) and CBDA (cannabidiolic acid) are chemicals uniquely produced by *Cannabis* plants. When decarboxylated to THC and CBD, these molecules bind to endocannabinoid receptors in the nervous systems of vertebrates and elicit a broad range of neurological effects in humans (*1*). Cannabinoid receptor types CB1 and CB2 preferentially bind THC and CBD, respectively, with CB1 being among the most abundant post-synaptic neuron receptor in the human brain whereas CB2 is more prevalent in the peripheral nervous system (*2-7*). Archeological and forensic evidence suggests that the psychoactivity of THC played a role in early domestication (*8-9*) and in selective breeding to increase marijuana potency during the late 20th century (*2*). Current explanations for the evolution of cannabinoid content focus on the duplication and divergence of *cannabinoid synthase* gene loci (*11-13*).

Domesticated *Cannabis* is divided into two major classes of cultivars: hemp and marijuana. Hemp, cultivated as a source of fiber, oil, and confectionary seed, produces modest amounts of CBDA and minimal THCA. Marijuana produces mostly THCA and much greater overall quantities of cannabinoids than hemp. Recent interest in CBD has led to the emergence of a new class of cultivars similar to marijuana. Like marijuana, these cultivars are generally short, highly branched plants with massive female inflorescences containing a high density of glandular trichomes and elevated cannabinoid content. Unlike marijuana, the predominant cannabinoid produced by these cultivars is CBDA. A principal component analysis of single nucleotide polymorphisms (SNPs) segregating in a diverse sample of *Cannabis* genotypes indicates that the THCA/CBDA ratio is associated with a major axis of population genetic differentiation (Fig. 1a). Hemp and marijuana cultivars are separated in the first principal component while the second component describes a continuum between naturalized populations and domestic cultivars. Estimated genetic divergence between population pairs (*Fst*_marijuana-naturalized_ = 0.128, *Fst*_hemp-naturalized_ = 0.147, and *Fst*_marijuana-hemp_ = 0.229) reflect a history of independent breeding trajectories with little gene flow between domesticated populations selected for divergent traits. However, economic incentives and regulatory policies that favored potent marijuana and non-intoxicating hemp in the past have shifted recently and plant breeders responded with targeted introgression.

**Fig. 1.**
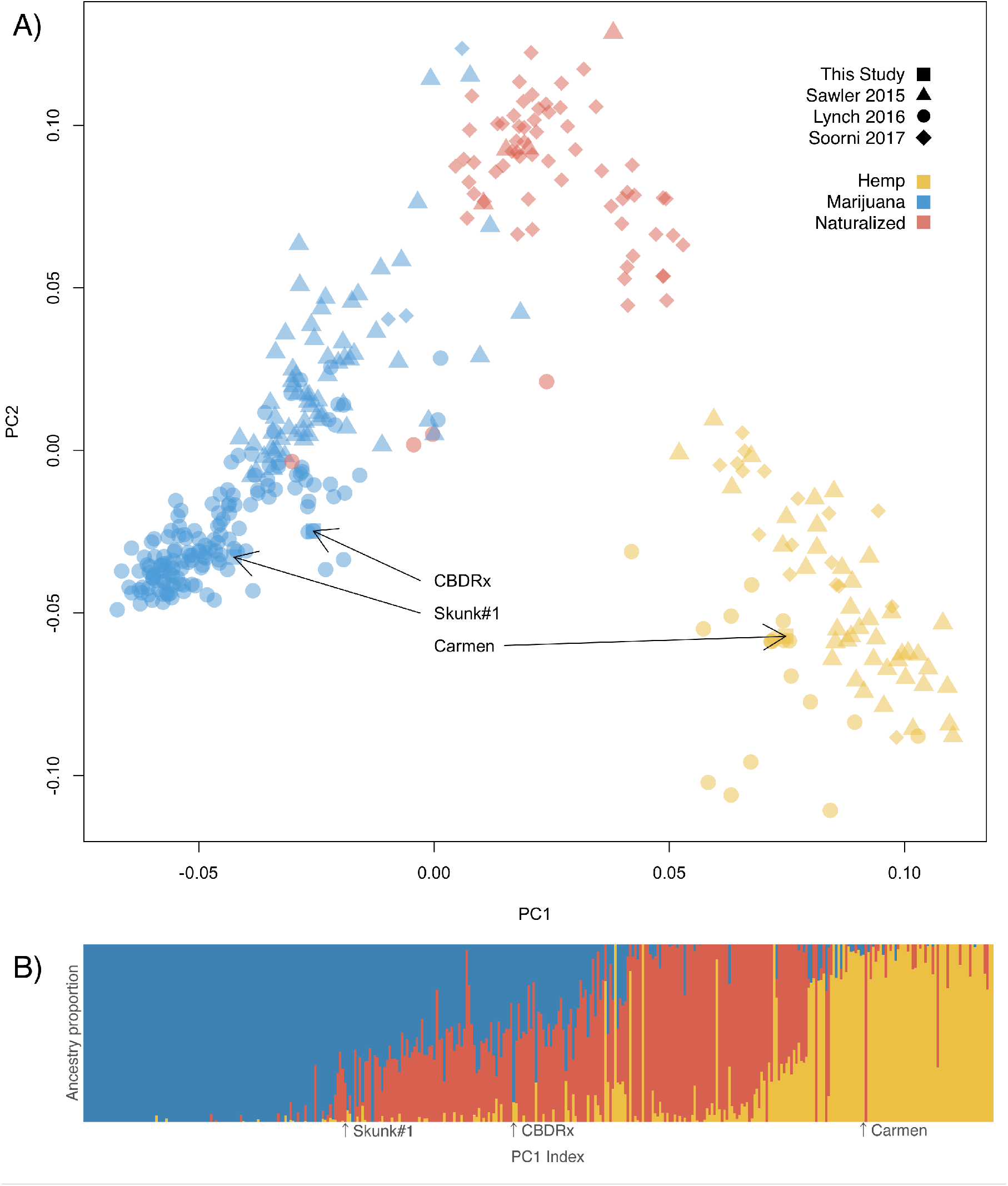
Marijuana and hemp are distinct populations of domesticated *Cannabis*. Population genetic structure of *Cannabis* inferred from 2,051 SNPs and 367 accessions delineates hemp, marijuana, and naturalized populations. The domesticated populations are both more closely related to naturalized populations than to each other. This reflects independent breeding trajectories with little gene flow between domesticated populations selected for divergent traits. Individuals were filtered to exclude relatives closer than the 5th degree. SNPs were filtered to reduce linkage disequilibrium and remove sites failing a chi-squared test for Hardy-Weinberg Equilibrium. (**A**) Principal components analysis (PCA) of the genotype matrix, integrating new data (plotted as squares) with previous population surveys (plotted as circles, triangles, or diamonds to indicate data source). Clusters were determined from k-means and are named according to a simple classification of hemp cultivars (yellow), marijuana cultivars (blue), and naturalized individuals (*10*). PC1 divides hemp and marijuana populations. PC2 describes the domestication continuum. The position of focal individuals with whole genomes sequenced in this study are indicated with arrows. Carmen is an industrial fiber hemp cultivar. Skunk#1 is an intoxicating marijuana cultivar. CBDRx has a predominantly marijuana-like genome, but is non-intoxicating. (**B**) Individuals are modeled with admixed genomes of idealized donor populations rather than being discreetly categorized. ADMIXTURE plot indicting ancestry contributions at k=3. Colors are consistent with populations defined for k-means classification on the PCA. Individuals are ordered left to right according to their position along the first principal component with their estimated ancestry proportions indicated by proportion of color contributing to the vertical segment. Focal individuals are indicated with arrows. The Skunk#1 genome ancestry is estimated to be 78% marijuana and 22% naturalized. The Carmen genome ancestry is estimated to be 94% hemp and 6% marijuana. The CBDRx genome is estimated to be 89% marijuana and 11% hemp.

The enzymes *THCA* and *CBDA synthase* (hereafter *THCAS* and *CBDAS*) compete for a common precursor (cannabigerolic acid or CBGA) and have been implicated in alternative explanations for the THCA/CBDA ratio. Some researchers focus on the role of sequence variation among *THCAS* gene copies (*3*), (*4*), while others (*5*) argue that the presence of a nonfunctional *CBDAS* allele in the homozygous state alters the cannabinoid ratio in favor of THCA. The public release of six *Cannabis* genomes, two of which were sequenced with long read technology, points to significant copy number variation among *synthase* genes across cultivars and yet their genomic structure has remained elusive (Table S1). The complexity of the *Cannabis* genome has also frustrated attempts to assemble complete chromosomes from thousands of contigs (Table S1), hindering the study of associations between *cannabinoid synthase* genes and drug potency.

In order to resolve the chromosomes of *Cannabis* and understand associations between *cannabinoid synthase* loci and cannabinoid content, we sequenced 100 whole genomes using a mixture of short and long read technologies. We sequenced near-isogenic marijuana (Skunk#1) and hemp (Carmen), an F1 hybrid, and 96 recombinant F2 individuals to construct an ultra-high-density genetic map and identify quantitative trait loci (QTL). We also used the genetic map to resolve the 10 *Cannabis* chromosomes (Fig. 2c) of the F1 and a high-CBDA cultivar (CBDRx) that were sequenced with long reads. Both genomes have higher contig contiguity than currently available *Cannabis* genomes (Table S1). These assemblies enabled us to completely resolve the *cannabinoid synthase* genes to three linked regions between 25-33 Mbp on CBDRx chromosome 9 (Fig. 2). The three regions are located on large contigs and contain 13 *synthase* gene copies. All but a single copy (located at 30Mbp) were found in two clusters of tandem arrays, consisting of seven (at 25 Mbp) and five copies each (at 29 Mbp). Each region has a single complete *synthase* coding sequence (Fig. 3d,e), with the two arrays having additional copies that are either incomplete or containing stop codons. All of the *cannabinoid synthase* loci are located in a highly repetitive pericentromeric region with suppressed recombination, and are linked in genetic and physical space (Fig. 2). The genomic context of these genes suggests distinct mechanisms by which copy number might evolve and differ among cultivars.

**Fig. 2.**
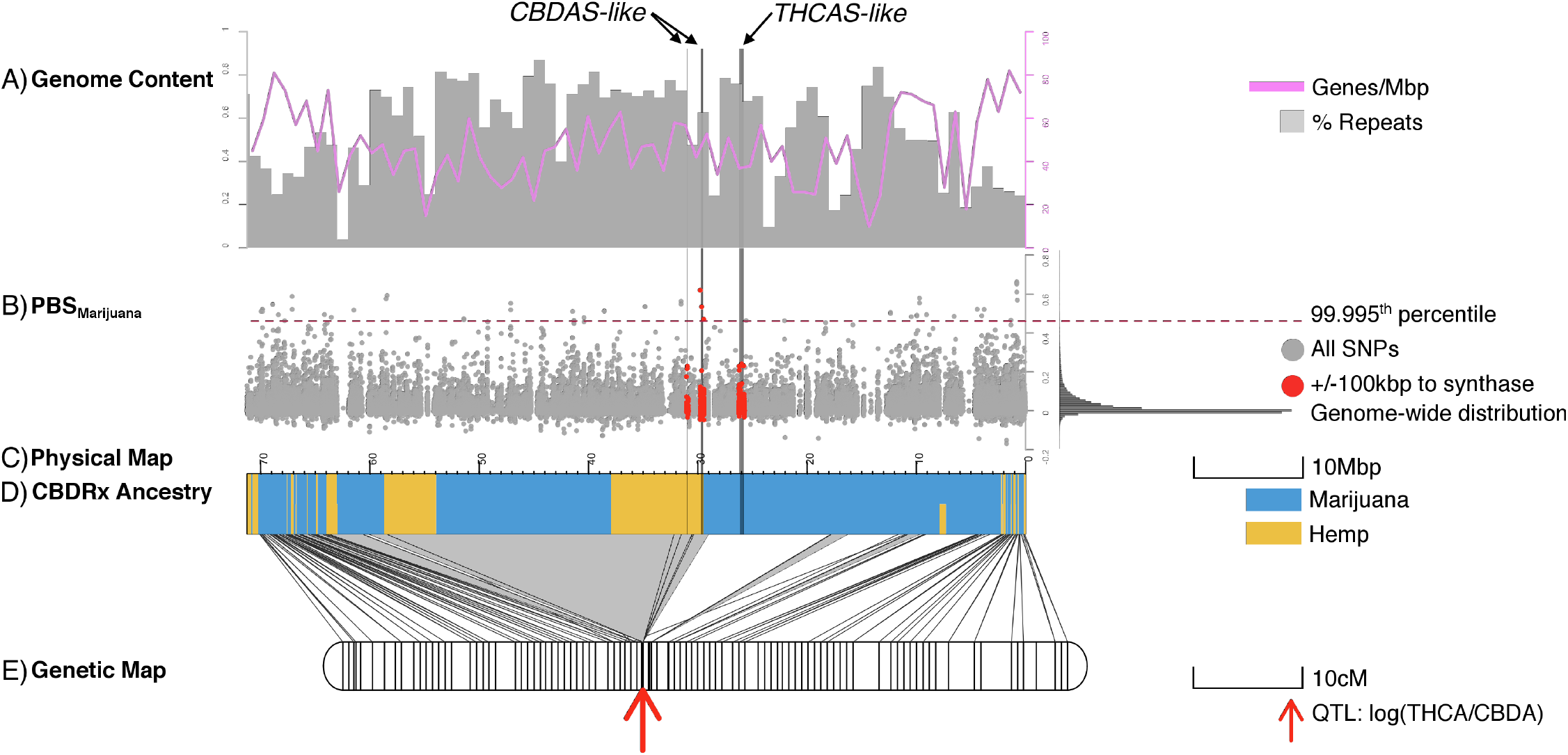
Genes responsible for chemotype on chromosome 9 are under selection in marijuana populations and have been targets for introgression by breeders. The locations of three cannabinoid synthase gene clusters are indicated by vertical lines transecting panels. Note that physical and genetic map coordinates are right-to-left. (**A**) Genes (pink lines) and percent repeat content (grey bars) in 1Mbp windows across the chromosome, (**B**) Manhattan plot of the population branch statistic (PBS), which is an *Fst*-based three-population test with extreme values suggesting lineage-specific evolutionary processes. The values for the marijuana branch are displayed here in grey dots across chromosome 9 with a histogram of the genome-wide distribution on the right. The 99.995^th^ percentile of the distribution is indicated with a dashed red line and values at SNPs within 100kbp of a cannabinoid synthase gene are indicated with red dots. We observe extreme values near *CBDAS* (but not *THCAS*), which is consistent with selection for nonfunctional *CBDAS* alleles in marijuana. (**C**) Painted ancestry of chromosome 9 in CBDRx with genomic segments derived from hemp in yellow and genomic segments derived from marijuana in blue. This analysis suggests a functional *CBDAS* allele from hemp was introgressed into a marijuana genome background to render the cultivar nonintoxicating. Ancestry blocks of CBDRx were called with AncestryHMM at SNPs separated by at least 0.3 cM and having high marijuana-hemp *Fst*. The genome-wide ancestry proportions of CBDRx were 89% marijuana and 11%. (**D**) The genetic map was anchored to the physical map using 211,106 markers segregating in an F2 mapping population. Lines connecting the genetic and physical maps indicate the positions of markers in physical and genetic space here. Physically consecutive markers with the same cM position have been consolidated to grey triangles. Grey triangles with the greatest area indicate regions of the genome with the least recombination. (**E**) A red arrow marks the position in the genetic map of the only QTL associated with the THC/CBD chemotype. This trait is perfectly correlated with the physical position of *CBDAS* and colocated with a genomic segment introgressed from hemp in the CBDRx genome. The total length of the genetic map is 818.6 cM, with a mean distance of 0.66 cM between observed crossovers.

The resolution of the three *cannabinoid synthase* regions with long read sequencing and correction-free assembly also provides insight into why the *THCAS* and *CBDAS* gene regions did not assemble previously (*4*) (Table S1). Each region is riddled with highly abundant transposable element sequences (Fig. 3e) and the two *synthase* clusters are comprised of 31-45 kb tandem repeats nested between Long Terminal Repeat (LTR) retrotransposons (Fig. 3a-c). The LTR (LTR08) associated with the *CBDAS* copies at 29 Mbp is predominantly restricted to this locus in the genome, and only small fragments of similar sequence were found on other chromosomes. In contrast, the LTR (LTR01) associated with *THCAS* repeats at 26 Mbp is found in high abundance over the entire genome and flanks the 29 Mbp cluster, suggesting that it may have played a role in the movement of the *CBDAS* cluster (Fig. 3). The fact that the LTR08 is specific to the *CBDAS* cassette in the genome further suggests it could be of distinct origin relative to the *THCAS* cassette.

**Fig. 3.**
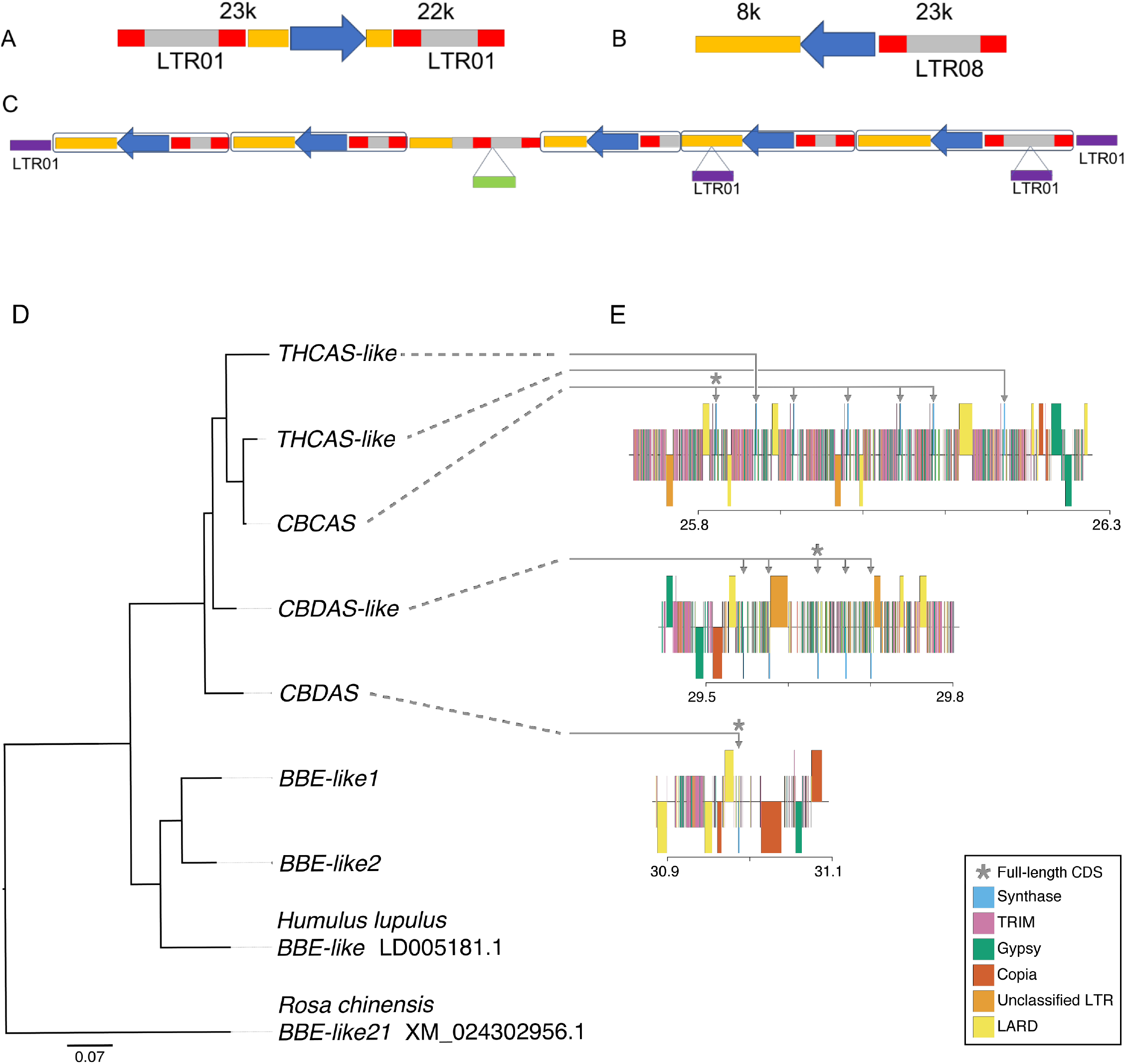
*Cannabinoid synthase* genes are located in tandemly repeated cassettes. Genes (blue) are clustered among long terminal repeats (LTR) colored as follows: LTR ends (*10*) LTR body (grey), unclassified LTR (orange), LTR01 remnants (purple), and an unclassified LTR fragment (green). The *synthase* gene cluster at 26Mbp includes seven copies of a cassette (A) ranging 38-84 kb in length and flanked by a pair of LTR01. *Synthase* genes at 29 Mbp are located in a different cassette (B) ranging 28-57 kb in length and having a single LTR08 upstream. (C) The entire 29 Mbp *synthase* gene cluster is flanked by LTR01 and the third cassette is interrupted by an LTR01 remnant. (D) CBDRx *cannabinoid synthase* gene tree rooted with closely related berberine bridge enzyme (BBE-like) sequences from rose (Rosa), hops (*Humulus*) and CBDRx. CBDRx sequences >97% similar are collapsed at the tips of the tree. (E) Functionally annotated maps of the *cannabinoid synthase* gene clusters in CBDRx. Genes in each of the three regions identified in Fig 2 are located in highly repetitive regions that include terminal repeat retrotransposons in miniature (TRIM), large retrotransposon derivatives (LARD), Gypsy, Copia and other unclassified long terminal repeats (LTR).

Coverage analysis confirmed that we identified 100% of the synthase gene copies in the CBDrx assembly (Table S3). In contrast, we identified 43 of 45 gene copies in the F1 assembly. These were resolved to either Carmen (22 copies) or Skunk #1 (23 copies) haplotypes. Most copies in the F1 assembly were solitary on short contigs, while one contig had three cassettes and seven contigs had two cassettes. Contigs bearing multiple cassettes confirmed the *synthase*-LTR tandem repeat structure. According to small size they could be not completely assembled, as was observed in previous assemblies like the Purple Kush genome where only 16% (5/30) of synthase homologs were assembled, all on short contigs (Table S3). That each *cannabinoid synthase* homolog within a tandem array shares the same promoter sequence suggests that variation in copy number within a gene cluster might have arisen by illegitimate recombination. However, another attractive model based on the architecture of the *synthase*-LTR tandem repeats is that breeding has selected for the activation and movement of *synthase*-LTR cassettes (*6*).

It is known from other systems that increases in copy number of biosynthetic gene clusters can elevate secondary metabolite production (*7*). Variation among *Cannabis* cultivars in the multiplicity of *cannabinoid synthase* loci (Table S2) encourages speculation that gene copy number might play a role in determining overall cannabinoid content. However, none of the five separate QTL we identified for total cannabinoid content (potency), were associated with *cannabinoid synthase* gene clusters (Fig. 4). For example, the strongest QTL for potency, accounting for 17% of variation in cannabinoid quantity, was located on chromosome 3 rather than chromosome 9. This suggests that traits and/or gene regulatory elements not linked to the *cannabinoid synthase* gene clusters affect cannabinoid quantity to a greater extent than the *synthases* themselves.

**Fig. 4.**
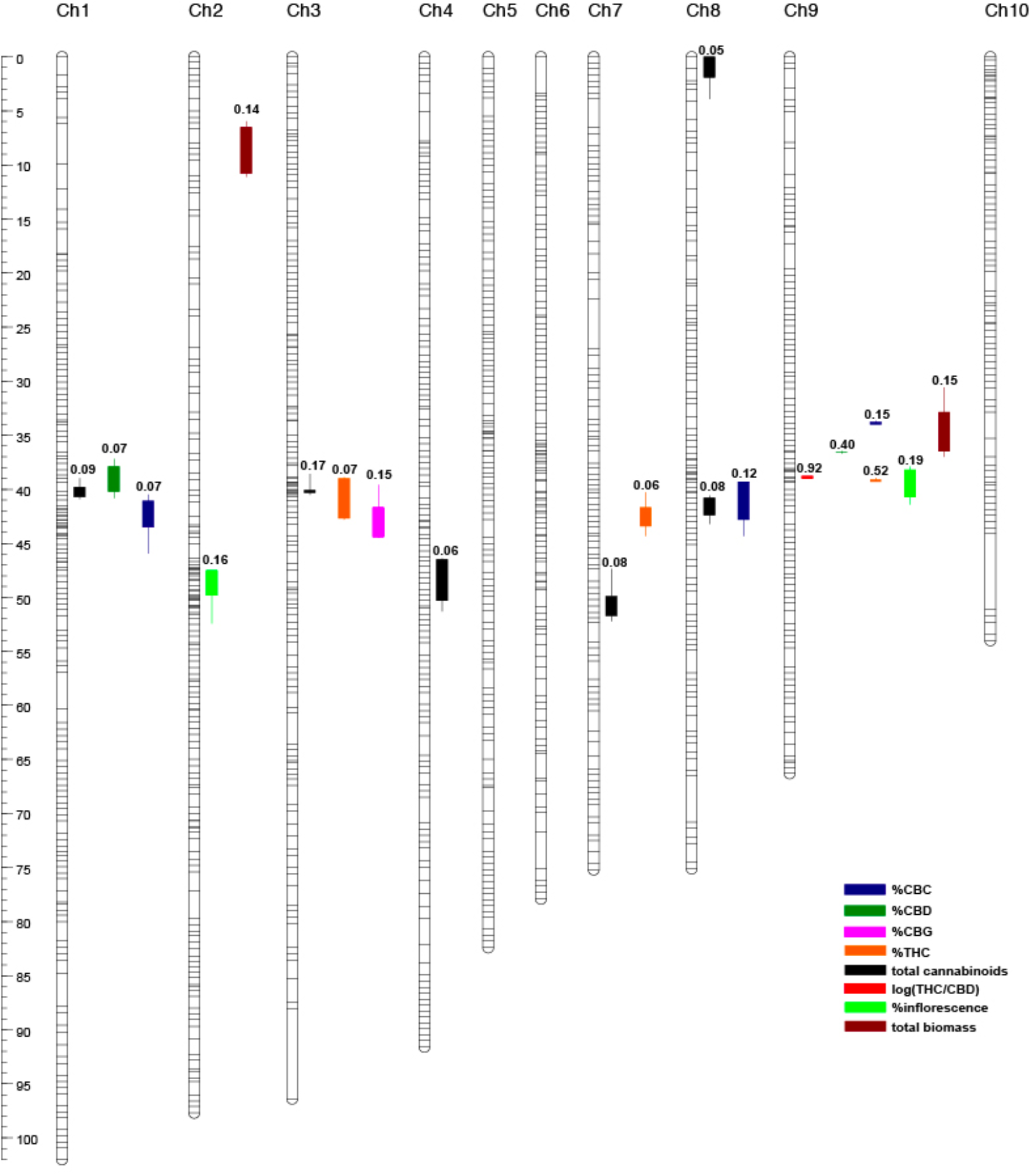
Composite genetic linkage and quantitative trait locus (QTL) map derived from a marijuana (Skunk#1) x hemp (Carmen) experimental cross. Map comprises ten linkage groups constructed from 211,106 markers segregating in 1,175 patterns from Illumina-based WGS of 96 F2 female plants integrated with 60 markers (48 AFLP, 11 microsatellite, 1 Sanger-sequence marker) scored across a subset of 62 F2 female plants (*5*). Segregation patterns represented as horizontal hash marks on linkage group bars. Quantitative trait loci (QTL) for ten phenotypes detected by composite interval mapping (P < 0.05; 1000 permutations) scored over 96 F2 plants indicated as vertical bar and whisker (1-LOD and 2-LOD intervals, respectively) plots to the right of corresponding linkage groups. Partial R^2^ for additive and dominance effects indicated above QTL plots. Genetic distance (centimorgans) scale bar to left of panel.

The *CBDAS* loci in particular appear to have been subject to recent selection in marijuana as evidenced by the population branch statistic (PBS) (*8*) (Fig. 2b) and dN/dS ratios (*5*). Contrary to the hypotheses of Onofri et al (*3*), these findings suggest that divergence at *CBDAS* loci rather than *THCAS* loci are primarily responsible for the THCA/CBDA ratio. We estimated genome-wide ancestry proportions of CBDRx to be 89% marijuana and 11% hemp. Most of the hemp-derived ancestry of CBDRx genome is found on only two chromosomes: 9 and 10. Notably, the genomic region associated with the QTL for log(THC/CBD), outlier branch lengths of the PBS genome scan for marijuana, as well as the identified *CBDAS* (but not *THCAS*), all lie within a shared segment with hemp ancestry. The *CBDAS* genes located at 29-31 Mbp are also nested in a region of CBDRx chromosome 9 with hemp ancestry whereas the *THCAS* tandem array is located in a region of marijuana ancestry. This pattern is consistent with the hypothesis that a predominantly *CBDA* cannabinoid profile is the result of introgression of hemp-like alleles into a marijuana genetic background to elevate *CBDA* production. That approximately 20% of chromosome 9 is hemp derived and tightly aligned with the QTL for the THCA/CBDA ratio provides further support for admixture combined with artificial selection resulting in new types of *Cannabis*, such as CBDRx, that present unprecedented combinations of phenotypic traits (*9*).

Here we generate the first chromosome scale assembly of the highly complex *Cannabis* genome, which required ultra-long nanopore sequencing reads and correction-free assembly to resolve the LTR-nested structure of the *THCA* and *CBDA synthase* tandem repeats on chromosome 9, two of which are traced back to hemp introgressions explaining the origin of high-CBDA cultivars. The architecture of the *synthase* loci suggests potential mechanisms for copy number variation, and strategies to manipulate these loci to improve cultivars. However, QTL results suggest that there are additional loci controlling potency in need of further investigation. After decades of regulation as a controlled substance, economic trends, recent changes in law, and the chromosome assembly presented here can accelerate the study of a plant that has co-evolved with human culture since the origins of agriculture.

### Data and materials availability

The *Cannabis* CBDRx and F1 genome and annotation are deposited at the European Nucleotide Archive under study PRJEB29284.

## Materials and Methods

### Plant material

CRBRx, a high-CBDA cultivar (15% CBDA and 0.3% THCA), was grown indoors in Colorado. Plants were grown in a compost enriched soil. CBDRx plants were grown indoors at 20-25C and 55-70% humidity under a mixture of fluorescent T-5 lamps and 1100W High Pressure Sodium Lamps manufactured by PL Lights. We made clonal cuttings approximately 10cm in height that included stems and leaves. These were immediately transferred to 42mm coconut coir plugs for rooting, then a coconut and perlite blend once roots were observed, where they remained for 40 days. Plants were then transferred to soil in 10cm pots for for 4 weeks of vegetative growth. Rooting and vegetative growth conditions included an 18:6 hour light:dark cycle and water as needed. Plants were transferred to 20L pots 10 weeks of flowering conditions using a 12:12 our light:dark cycle. Plants were fertilized with a micronutrient blend certified by the Organic Materials Review Institute plus biochar. Under flowering conditions, plants were watered every 7-9 days. A single plant (CBDRx:18:580) was chosen while in the vegetative phase and recently emerged leaves were collected for DNA purification.

The genetic background and cultivation of the mapping population over which the linkage and QTL mapping are reported has been previously described (*5*). In brief, parental marijuana (Skunk#1) and hemp (Carmen) lines were sibling crossed for five generations to increase homozygosity. A single fifth-generation Skunk#1 female was fertilized with pollen from a single fifth-generation Carmen male. From the resulting seed, a single genetically female F1 (CO9) plant was isolated and vegetatively cloned. Stamen development was induced in mature pistillate CO9 clones via treatment with colloidal silver, resulting in monoecious plants. CO9 clones were fertilized with pollen from CO9 clones to produce an F2 seed generation. Female F2 plants were grown from seed to flowering maturity for 12 weeks under conditions previously described (*5*). Mature flowers of the parents and F2 plants were collected at harvest and dried for subsequent DNA purification.

A single male F1 (CO11) plant full-sibling to the F1 (CO9) from which the mapping population descended was grown from seed under vegetative light (16h light: 8hr dark) and high nitrogen nutrient conditions equivalent to the initial four weeks of growth used for the mapping population (*5*) except that LED lighting (Valoya R150-NS1; Valoya Oy, Helsinki, Finland) was used. Fresh recently emerged leaves were collected from this plant for high molecular weight DNA purification.

### Cannabinoid analysis

Cannabinoid analysis by GC was as described in Weiblen et al., (*5*).

### Agronomic trait phenotyping

F2 plants were grown for four weeks under vegetative conditions followed by eight weeks under flowering conditions (*5*). After twelve weeks of growth, plant height was measured from the base of the primary stem to shoot apex after which plants were harvested at the stem base and dried for three weeks at ambient conditions. Dried plant tissue fractions (stems, leaves, inflorescences) were weighed and percent mass of each fraction was calculated relative to total harvested mass.

### Illumina sequencing

We extracted DNA from 15-20 mg of dried flowers from each of Skunk#1, Carmen, and 96 F2 individuals using a microfuge-scale CTAB-buffer/organic extraction protocol (adapted from (*11*). Isolated DNAs were quantified using the PicoGreen dsDNA assay kit (ThermoFisher), size-evaluated by Agilent TapeStation gDNA (Agilent, Santa Clara CA) and used as input for TruSeq DNA PCR-Free (Illumina, San Diego CA). All 96 PCR-free libraries from the F2 set were pooled on an equimolar basis using PicoGreen concentrations. Likewise, a second pool was created from the Skun#1 and Carmen libraries. We used quantitative PCR (qPCR) to assess functionality, which was approximately 25%. Each library pool was adjusted according to the qPCR results prior to sequencing. Libraries were sequenced on an Illumina HiSeq 2500 SBS V4 in 2×125bp read high-output mode (Illumina, San Diego CA) at the University of Minnesota Genomics Center. Raw reads were gently trimmed of low-quality bases and synthetic sequence using Trimmomatic (*12*).

For all Illumina data we trimmed reads of adapter sequence with Trimmomatic (*12*), aligned them to the reference assembly with BWA MEM (*13*), sorted and compressed the alignments with Samtools (*14*), and marked duplicates with Picard tools (*15*).

### PacBio sequencing

Genomic DNA of the F1 was obtained from fresh young leaf tissue using a modified CTAB/organic extraction protocol (adapted from (*11*) in which the extraction buffer was supplemented with antioxidants (0.5% sodium diethyldithiocarbamate, 10mM sodium metabisulfite), and a DNAse inhibitor (200mM L-lysine). Precipitated DNAs were collected using a glass hook, rinsed with ethanol, and resuspended in deionized water. Genomic DNA was quantified using the PicoGreen dsDNA assay kit (ThermoFisher), size-evaluated by Agilent TapeStation gDNA and pulsed-field gel electrophoresis (PFGE), diluted to 50 ng/uL, and sheared via 20 passes through a 26G blunt needle. Shears were evaluated using PFGE. Approximately 15 μg of sheared and concentrated DNA was used as input into library prep using the SMRTbell Template Prep Kit 1.0 using a protocol for >30kb libraries (101-181-000 Version 05). The resulting library was size-selected with a 20 kb high-pass protocol using the PippinHT, and an additional DNA Damage Repair was performed to generate the final library. Sequencing was performed via diffusion loading with Sequel Binding Kit 2.0 and a mixture of Sequel Sequencing Kits 2.0 and 2.1.

### Nanopore sequencing

Leaf material from the inbred CBDrx line was flash frozen in liquid nitrogen. 5 g of flash frozen leaf tissue was ground in liquid nitrogen and extracted with 20 mL CTAB/Carlson lysis buffer (100mM Tris-HCl, 2% CTAB, 1.4M NaCl, 20mM EDTA, pH 8.0) containing 20μg/mL proteinase K for 20 minutes at 55°`C`. The DNA was purified by addition of 0.5x volume chloroform, which was mixed by inversion and centrifuged for 30 min at 3000 RCF, and followed by a 1x volume 1:1 phenol: [24:1 chloroform:isoamyl alcohol] extraction. The DNA was further purified by ethanol precipitation (1/10 volume 3 M sodium acetate pH 5.3, 2.5 volumes 100% ethanol) for 30 minutes on ice. The resulting pellet was washed with freshly-prepared ice-cold 70% ethanol, dried, and resuspended in 350 μL 1x TE buffer (10 mM Tris-HCl, 1 mM EDTA, pH 8.0) with 5 μL RNase A (Qiagen, Hilden) at 37°C for 30 min, followed by incubation at 4°C overnight. The RNase A was removed by double extraction with 24:1 chloroform:isoamyl alcohol, centrifuging at 22,600x*g* for 20 minutes at 4°C each time. An ethanol precipitation was performed as before for 3 hours at 4°C. The pellet was washed as before and resuspended overnight in 350 μL 1x TE.

Genomic DNA sample was further purified for Oxford Nanopore (ONT) sequencing with the Zymo Genomic DNA Clean and Concentrator-10 column (Zymo Research, Irvine, CA). The purified DNA was then prepared for sequencing following the protocol in the genomic sequencing kit SQK-LSK108 (ONT, Oxford, UK). Briefly, approximately 1 μg of purified DNA was repaired with NEBNext FFPE Repair Mix for 60 min at 20°C. The DNA was purified with 0.5X Ampure XP beads (Beckman Coulter). The repaired DNA was End Prepped with NEBNExt Ultra II End-repair/dA tail module including 1 μl of DNA CS (ONT, Oxford, UK) and purified with 0.5X Ampure XP beads. Adapter mix (ONT, Oxford, UK) was added to the purified DNA along with Blunt/TA Ligase Master Mix (NEB, Beverly, MA) and incubated at 20°C for 30 min followed by 10 min at 65°C. Ampure XP beads and ABB wash buffer (ONT, Oxford, UK) were used to purify the library molecules and they were recovered in Elution Buffer (ONT, Oxford, UK). Purified library was combined with RBF (ONT, Oxford, UK) and Library Loading Beads (ONT, Oxford, UK) and loaded onto a primed R9.4 Spot-On Flow cell. Sequencing was performed with a MinION Mk1B sequencer running for 48 hrs. Resulting FAST5 files were base-called using the ONT Albacore software using parameters for FLO-MIN106, and SQK-LSK108 library type.

### Full length cDNA sequencing with Oxford Nanopore

Fresh CBDrx leaf tissue was flash frozen in liquid nitrogen and ground to a fine powder using a mortar and pestle. RNA was extracted from the powder using the Qiagen Plant RNeasy Plant Mini Kit (Qiagen, Netherlands). RNA quality was assesed using a bioanalyzer. High quality RNA was used to generate full length cDNA using the cDNA-PCR Sequencing Kit (SQK-PCS108, Oxford Nanopore Technologies, Oxford, UK). Resulting libraries were sequenced on the Oxford Nanopore GridION sequencer (Oxford Nanopore Technologies, Oxford, UK) for 48 hrs.

### Nanopore genome assembly

A total of 27 Gb of Oxford Nanopore sequence was generated on the MinION ONT platform. The resulting raw reads in fastq format were aligned (overlap) with minimap and an assembly graph (layout) was generated with miniasm2 (*16*). The resulting graph was inspected using Bandage (*17*). A consensus sequence was generated by mapping reads to the assembly with minimap2, and then Racon (*18*) three times. Finally, the assembly was polished with pilon (*19*) three times using the Illumina paired-end 2×100 bp sequence; the Illumina reads were mapped to the consensus assembly using BWA (*13*). All assembly steps were carried out on a machine with 231 Gb RAM and 56 CPU.

### Assembly Results

We sequenced CBDRx to a 34x coverage using long read Oxford Nanopore Technology (ONT) and the F1 to a 5x circular consensus coverage using PacBio Single Molecule Real-Time (SMRT) long read sequencing for the purpose of comparing THCAS and CBDAS variation in our mapping population to CBDRx. Both genomes were assembled using a correction-less assembly pipeline that consisted of an overlap (minimap2), layout (miniasm2) consensus (racon), followed by a polishing step (pilon) using the 64x Illumina 2×100 bp paired end reads (*20*). The resulting CBDRx assembly was 746 Mbp in 1,986 contigs with an N50 length of 742 kb and the longest contig 4.5 Mbp, while the F1 assembly was 1,389 Mbp in 12,204 contigs with an N50 length of 172 kb and the longest contig 1.9 Mbp (Table S2). Both genomes have higher contig contiguity than the Cannabis genomes currently available (Table S1), and form the basis for a complete chromosome assembly.

### Genetic Linkage Map

Our core mapping population is made up of F2s germinated from seed collected from the CO9 clones. A pseudo F1 dataset was constructed by concatenating all F2 reads followed by random subsampling to a target genomic coverage of 100x. The pseudo F1 and parental reads were independently error corrected using k-mer histograms with k=25 with AllpathsLG (*21*). A de Bruijn graph was constructed from the error corrected pseudo F1 reads using McCortex assembler at k=19 (*22*). This program is unique in that genome assembly and variant discovery are performed simultaneously - reads are assembled, but the paths through “bubbles,” i.e. regions of the graph that diverge and rejoin are retained as variants. The bubble read coverage distribution is used to classify bubbles as repeats, homologous alleles, or errors. Parental reads and F2 reads were threaded through the graph independently. F2s were genotyped at variant sites at which Carmen and Skunk#1 were fixed for alternate alleles. Genotypes were updated via imputation using a sliding-window hidden markov model using LB-Impute (*23*) leveraging variant coverage information and physical linkage within a window of width 10 variants.

Segregation patterns of genotypes containing no missing data across the population that appeared at least ten times were selected for use as map markers. Markers exhibiting segregation distortion by Chi^2 test were low in number and are retained in the map (~10% of markers). Linkage groups and marker order were inferred using the ant colony optimization in AntMap (*24*) solution to the traveling salesman path. Recombinations were counted directly and divided by the number of gametes in the population (192) to infer genetic distance between adjacent markers and summed consecutively in linear order to give map position on a linkage group.

### Linkage mapping

Markers obtained from a high-density map made using Illumina data built using AntMap for 96 F2 individuals were used to produce a composite map built by adding markers from Weiblen (*5*) using JOINMAP 4.1 (Wageningen, the Netherlands). Linkage groups were assembled from independent log-of-odds scores (LOD) based on G-tests for independence of two-way contingency tables. Linkage groups with LOD > 3.0 and containing four or more markers were used to construct a linkage map using the Kosambi (*25*) function. The high-density composite linkage map comprises ten linkage groups, 1,235 total segregation patterns, a map distance of 818.6 cM and a mean intermarker distance of 0.66 cM.

### QTL analysis

Cannabinoid profiles and biomass traits of the same 96 F2 individuals were analyzed with respect to the composite linkage map using Windows QTL CARTOGRAPHER v.2.5_011 (*26*); WinQTLCart). Composite interval mapping was used to estimate LOD over a walk speed of 1.0 cM and significant associations between traits and linkage groups were identified using an experiment-wise (P = 0.05) LOD threshold estimated in WinQTLCart using 1000 permutations. Results were plotted with MAPCHART 2.32 (Wageningen, the Netherlands).

### Pseudomolecule generation

The genome assembly was evaluated for library contaminants using Blobtools (*27*) and the NCBI non-redundant database. Contigs with good evidence as derived from outside viridiplantae were removed. We aligned the genetic map bubbles to the CBDRx contigs with BWA (*13*). Contigs were deemed chimeric if they mapped to different linkage groups or more than 10 centimorgans away from each other and broken at the longest repeat between genetically mapped regions. An initial set of rough pseudomolecules were constructed by assigning contigs to linkage groups, ordering contigs by mean centimorgan (*28*), and orienting by cM position on either end. The F2 population was genotyped again via alignment to the rough pseudomolecules followed by LB-Impute. Population segregation patterns from this second round of genotyping were used to further saturate the genetic map if they increased map density without increasing the map length (*29*). Contigs were partitioned by linkage group and scaffolded with the Hi-C library using three iterations of Salsa (*30*). Allmaps was used to generate the final contig order and orientation with the template genetic map positions, second round genetic map positions, and Salsa contig positions as input. The pseudomolecules were further polished with an additional ten iterations of Racon followed by an additional ten iterations of Pilon. After chromosome-wide scaffolding and gap filling, 841 contiguous sequences spanning 714,498,588 bp were anchored to nuclear pseudomolecules. We genotyped the genetic mapping population for a third time against the CBDRx reference and visually inspected the segregation patterns for misorderings. We found most contigs to be largely collinear in genetic and physical space. We observed zero recombinanants on a minority of contigs and were unable to resolve their relative order and orientation. This was the case for two of the three synthase-bearing contigs on chromosome 9. For these, we manually reordered the synthase-bearing contigs to be physically adjacent, as we could not find evidence supporting an alternative arrangement and such an arrangement is most parsimonious with study-wide results.

### Repeat and gene prediction and annotation

Full length LTRs were predicted using LTRfinder using the standard settings and 1 mismatch (*31*). The resulting full length LTRs were used to mask the genome using repeat masker (*32*). Four full-length cDNA nanopore read libraries were aligned to the reference with minimap2 (*33*) before and after error correction by Canu (*34*) of colocated batches. 142 RNAseq libraries found on the Sequence Read Archive were aligned to the reference with GSnap (*35*) and assembled into transcripts with Stringtie (*36*). 4 high-coverage RNASeq libraries were assembled using Trinity (*28*) in both de-novo and reference-guided modes. Contaminate sequence was removed using Seqclean (*15*). The full-length cDNAs, Stringtie assembly, and Trinity transcripts were assembled into gene models with the Program to Assemble Spliced Alignments (*37*). Additional transcriptome assemblies from *Humulus lupulus* (*38*) and *Cannabis* were aligned to the reference with GMap. Genes were predicted ab initio using Augustus (*39*). Non redundant RefSeq proteins (*40*) for viridiplantae were clustered at 90% identity with CD-HIT (*41*). Representative sequences for each cluster were aligned to the reference genome using Diamond (*42*) —extra-sensitive. Pairwise hits were locally realigned with AAT (*43*) and Exonerate protein2genome. Repetitive sequence was identified using the set union of three programs: RepeatMasker, Tephra, and Red (*44*). EvidenceModeler was used to integrate all evidence for and against protein-coding genes. PASA was updated with these results.

### CBDRx chromosome assembly analysis

The CBDRx ONT-based contig assembly was further resolved into chromosomes using a genetic map derived from progeny of the F1. Using whole-genome-shotgun sequencing (WGS) we scored 96 F2 plants for 211,106 markers segregating in 1,235 high-confidence patterns resulting in ten linkage groups, which we then used to anchor the ONT-based contigs. The final chromosome-resolved assembly of CBDRx captured 90.8% of the gene space as predicted by Benchmarking Universal Single Copy Orthologs (BUSCO) (*45*). The CBDRx genes were predicted using a combination of *ab initio* and empirical data including full length cDNA sequenced using ONT long read sequencing, as described above. After masking 63% of the genome for repeats that were made up of 17,536 full length long terminal repeats (LTRs), 42,052 protein coding genes were predicted in the CBDrx assembly. We identified the 345-355 bp subtelomeric repeat that has been defined in *Humulus lupulus* (*46*), and the 224 bp centromeric repeat (*47*). That 17% of the reads mapped to the centromere repeat and 14% mapped to the subtelomeric repeat is consistent with their predicted size in the genome. These observations support the first complete *Cannabis* chromosome assembly and a framework for examining the genomic structure of the *THCAS* and *CBDAS* loci in association with quantitative traits.

### THCAS/CBDAS and coverage analysis

In addition to the gene prediction and annotation, and *CBDAS* and *THCAS* genes (AB292682 and AB057805 respectively) were used to search the final assembly. The *THCAS* and *CBDAS* gene sequence was blasted against the final CBDrx assembly to confirm their locations. We identified four locations on the genome with close hits to *synthases*:, chr9:26 Mbp, ch9:29 Mbp, ch9 31 Mbp, and the more-distantly related homologs at chr6:15 Mbp. To check that all of the genes were captured in the assembly a coverage analysis was performed. The Illumina reads were mapped to a single copy gene *GIGANTEA (GI*), one ribosomal cassette (18S-5S-26S) and the four version of the *synthase* genes. The results confirmed the 14 *synthase* genes and suggested 500-600 rDNA arrays, which is consistent with other genomes this size (Table S3). Coverage analysis in Carmen, Skunk#1 and Purple Kush (*4*) revealed 22, 24 and 30 *synthase* genes respectively.

### Comparative Genomics

Individuals with sequenced WGS libraries were genotyped with BCFtools. Genome-wide ancestry proportions at k=3 were estimated using ADMIXTURE (*48*). Individuals identified as having >99% ancestry were assigned to respective marijuana and hemp populations. A subset of segregating sites were selected for assigning ancestry tracts along chromosomes using a method intended to maximize informativeness and minimize linkage disequilibrium. Sites were ranked by Wright’s *Fst* (*49*). Genetic positions for all segregating sites were interpolated along a B-spline function fitted to the empirically observed positions in the mapping population with coefficients penalized to maintain monotonicity (*50*). For each chromosome, the site with the highest *Fst* value and lowest genetic position was the first selected. Decreasing by *Fst* through all segregating sites, additional sites were selected so long as they were at least 0.03 cM from any previously selected site. Ancestry tracts were assigned by AncestryHMM (*51*) assuming a single pulse from hemp to marijuana eight generations in the past.

We obtained previously published population data from the original authors and genotyped individuals against the CBDRx reference using our standard pipeline including sites with a quality score greater than 500. In order to understand neutral population structure, we used Plink and Plink2 to filter the genotype matrix to minimize structure originating from familial relatedness, artifactual patterns in occupancy, selection, and genetic linkage. We selected a single representative individual from groups with KING-robust kinship coefficient greater than 0.015625. We retained bi-allelic sites called in at least 80% of individuals, with a minor allele frequency greater than 1%, observed heterozygosity less than 60%. We removed sites failing an exact test for Hardy-Weinberg at p-value of 1e-20 with a mid-p adjustment (*52*). We eliminated individuals genotyped at less than 90% of sites. We thinned sites for linkage disequilibrium in sliding windows with a width of 50 SNPs, a slide of 5 SNPs, and a variance inflation factor threshold of 2. We used this plink-filtered genotype matrix for PCA and k-means clustering, as well as Admixture analysis at k=3. We used the Population Branch Statistic to scan the genome for sites undergoing population-specific processes. We assigned individuals to populations based on their k-means cluster membership and retained all sites with a quality score greater than 500 for this analysis. We calculated *Fst* (*53*) for the three population pairs using VCFtools. The PBS is three population test. For populations (a,b,c):

PBS_a = ((T_ab + T_ac − T_bc) / 2)
PBS_b = ((T_ab + T_bc − T_ac) / 2)
PBS_c = ((T_ac + T_bc − T_ab) / 2)
with
T_ab = −log(1 − Fst_ab)
T_ac = −log(1 − Fst_ac)
T_bc = −log(1 − Fst_bc)

## Acknowledgments

Veronica Tonnell contributed DNA isolation for the mapping population. We thank Matthew Gibbs and Jason Schwartz for support from Sunrise Genetics Inc. Timothy Gordon of Functional Remedies provided the high-CBDA plant. C.J.G. is grateful to Cristina M. Moya for insightful conversations on human-plant interactions as well as hosting his sabbatical at the Max Planck Institute of Evolutionary Anthropology.

## Author contributions

G.D.W. and C.J.G, and C.J.S. designed the study. G.D.W., J.P.W., and C.D. developed the mapping population and prepared materials for genomic analysis. T.P.M, S.G.P, and S.T.M sequenced the CBDRx genome and full length cDNA. C.J.G. and T.P.M assembled, annotated, and analyzed the genomes. C.J.G. integrated the maps and analyzed the populations. J.P.W and C.D. measured phenotypes and performed QTL analysis. C.J.G., C.D., J.P.W., C.J.S., T.P.M and G.D.W. wrote the manuscript.

---

**Table S1.** Sequenced Cannabis genomes fail to resolve the THCA/CBDA loci.

| Cultivar | GenBank | Research Group | Date | Contigs | Coverage | N50 | CBDA/THCA genes | Contigs | Method |
| --- | --- | --- | --- | --- | --- | --- | --- | --- | --- |
| Purple Kush | ASM23057 | University of Toronto | 10/18/11 | 135,164 | 130x | 16,377 | 5 | 5 | Illumina GA II; HiSeq |
| LA Confidential | ASM151000 | Courtage Life Sciences | 1/11/16 | 311,039 | 50x | 2,649 | 6 | 6 | Illumina 454 |
| Chemdog91 | Chemdog91_175268 | Courtage Life Sciences | 1/11/16 | 175,088 | 300x | 2,250 | 1 | 1 | Illumina GA II |
| Cannatonic | ASM186575 | Phylos Bioscience | 11/3/16 | 11,110 | 130x | 128,718 | 12 | 12 | PacBio |
| Pineapple Banana Bubba Kush | ASM209043 | Steep Hill Genetics/ CU Boulder, CGRI | 4/13/17 | 18,355 | 72x | 51,819 | 12 | 12 | PacBio |
| Finola | 003417725.1 | University of Toronto. Anandia | 8/22/18 | 10,878 | 97x | 116,560 | 10 | 10 | PacBio |

**Table S2.** CBDRx and F1 genome assembly statistics.

|  | CDBRx | F1 (Carmen x Skunk #1) |
| --- | --- | --- |
| Contig (#) | 1,986 | 12,202 |
| Largest contig (bp) | 4,475,621 | 1,889,784 |
| Total Length (bp) | 747,554,284 | 1,389,290,832 |
| GC (%) | 33 | 34 |
| N50 contig length (bp) | 742,283 | 172,909 |

**Table S3.** Coverage analysis using Illumina reads and the assembled CBDAs/THCAs genes.

|  | **CBDRx** | **Carmen** | **Skunk** | **Purple Kush** |
| --- | --- | --- | --- | --- |
|  | Nanopore;<br>Illumina | CD1_illumina;<br>Pacbio<br>(haplotype<br>resolved from<br>F1) | CF2_illumina;<br>Pacbio<br>(haplotype resolved from F1) | CanSat |
| CBDAs Chr09 29 Mb | 5 | 6 | 8 | 7 |
| CBDAs_Ch09 30 Mb | 1 | 2 | 4 | 7 |
| BBE-like Chr06 | 1 | 2 | 2 | 5 |
| THCAs_Ch09 25 Mb | 7 | 13 | 10 | 11 |
|  | 14 | 22 | 24 | 30 |
| F1 |  | 46 |  |  |
| Assembly | 14 | 43 |  | 5 |

**Table S4.** QTL composite interval mapping results of phenotypic traits.

| Trait | Chromosome# | Peak LOD | Peak LOD<br>cM position | 1-LOD L | 1-LOD R | 2-LOD L | 2-LOD R |
| --- | --- | --- | --- | --- | --- | --- | --- |
| %CBD | 1 | 6 | 39.43 | 37.91 | 40.15 | 37.21 | 40.79 |
| %CBD | 9 | 20.34 | 36.57 | 29.49 | 51.67 | 36.53 | 36.72 |
| %CBC | 1 | 5.36 | 41.46 | 41.07 | 43.49 | 40.47 | 45.92 |
| %CBC | 8 | 7.8 | 41.32 | 39.33 | 42.84 | 39.3 | 44.26 |
| %CBC | 9 | 9.45 | 33.88 | 33.81 | 33.95 | 33.74 | 34.02 |
| %THC | 3 | 5.19 | 41.65 | 38.95 | 42.72 | 38.93 | 42.83 |
| %THC | 7 | 4.6 | 43 | 41.71 | 43.38 | 40.31 | 44.28 |
| %THC | 9 | 25.08 | 39.19 | 39.09 | 39.26 | 38.99 | 39.33 |
| %CBG | 3 | 6.56 | 44.31 | 41.74 | 44.39 | 39.59 | 44.47 |
| %inflor | 2 | 4.96 | 48.39 | 47.47 | 49.83 | 47.45 | 52.41 |
| %inflor | 9 | 7.39 | 39.43 | 38.18 | 40.71 | 37.86 | 41.36 |
| totbio | 2 | 5.35 | 9.6 | 6.53 | 10.75 | 6 | 11.1 |
| totbio | 9 | 6.2 | 35.47 | 32.9 | 36.48 | 30.62 | 36.96 |
| totcann | 1 | 6.78 | 40.59 | 39.77 | 40.74 | 39.03 | 40.89 |
| totcann | 3 | 11.38 | 40.22 | 40.08 | 40.34 | 38.57 | 40.5 |
| totcann | 4 | 5.02 | 49.28 | 46.53 | 50.33 | 46.43 | 51.31 |
| totcann | 7 | 5.83 | 50.24 | 49.86 | 51.7 | 47.35 | 52.22 |
| totcann | 8 | 4.41 | 1.13 | 0.01 | 1.91 | 0.01 | 3.94 |
| totcann | 8 | 5.83 | 41.32 | 40.77 | 42.35 | 40.63 | 43.18 |
| logTHC/CBD | 9 | 60.23 | 38.89 | 38.82 | 38.95 | 38.75 | 39.03 |

